# Faster and better CRISPR guide RNA design with the Crackling method

**DOI:** 10.1101/2020.02.14.950261

**Authors:** Jacob Bradford, Timothy Chappell, Dimitri Perrin

## Abstract

The design of CRISPR-Cas9 guide RNAs is not trivial, and is a computationally demanding task. Design tools need to identify target sequences that will maximise the likelihood of obtaining the desired cut, whilst minimising off-target risk. There is a need for a tool that can meet both objectives while remaining practical to use on large genomes.

Here, we present Crackling, a new method that is more suitable for meeting these objectives. We test its performance on 12 genomes and on data from validation studies. Crackling maximises guide efficiency by combining multiple scoring approaches. On experimental data, the guides it selects are better than those selected by others. It also incorporates Inverted Signature Slice Lists (ISSL) for faster off-target scoring. ISSL provides a gain of an order of magnitude in speed, while preserving the same level of accuracy. Overall, this makes Crackling a faster and better method to design guide RNAs at scale.

Crackling is available at https://github.com/bmds-lab/Crackling under the Berkeley Software Distribution (BSD) 3-Clause license.

## Introduction

The design of this guide RNA is a crucial step for any CRISPR experiment. However, guide design is not trivial, as the efficiency and specificity of guides are crucial factors. Efficiency broadly refers to the guide correctly binding to the targeted region and to the endonuclease. This is influenced by a number of factors that depend both on the guide itself (e.g. secondary structure) and on the targeted region (e.g. chromatin accessibility). Specificity refers to the need for the guide RNA not to induce off-target modifications. This means not only that the targeted sequence must be unique, but also that closely related sequences (typically less than four mismatches) elsewhere in the genome must be carefully considered.

We recently benchmarked existing guide design tools [1]. Our two main findings were that:

1. When considering efficiency, for any given genomic sequence there is a limited overlap between the set of guides that each tool is producing.
2. Several tools have inadequate filtering on specificity, and when that is considered carefully, it tends to be a computationally expensive task.

The limited overlap between the tools can be exploited for guide selection: when a guide is recommended by multiple tools, there is a higher chance that it will actually be efficient. We explored consensus approaches in detail [2], and they provide better guides. However, it is not necessarily practical to run several tools. A recent review suggested to use a tool which offers more than one scoring algorithm and these exist, however, none have explored the possibility of combining the scores to achieve an overall gain in precision [3]. Our *Crackling* method directly integrates multiple scoring approaches and combines their result for improved precision when predicting guide efficacy.

Guide efficacy is critical, yet the success of a CRISPR experiment is also determined by the specificity of a guide RNA. A number of methods have been implemented to tackle specificity evaluation. One of the earliest methods, published in 2014, is Cas-OFFinder [4]. It provides a count of the number of mismatches between each candidate guide and off-target site, if the count is less than some threshold (e.g. four). The results can then be used to get a sense of the off-target risk for each candidate guide.

While identifying off-target sites that are within a fixed number of mismatches is useful, it is not necessarily enough. It has been experimentally shown that the position of the mismatches also matters [5]. A score for this exists; sometimes referred to as the Zhang score, from the corresponding author of that paper. To calculate the score, a candidate guide is compared against all potential CRISPR sites in the genome of interest. When a site is at most four mismatches away, a local score is calculated using this formula:

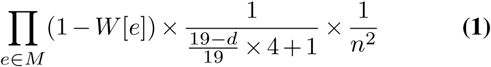

where *W* is an array of position-specific weights, *d* is the pairwise distance between mismatches, *n* is the number of mismatches between target and sequence, and *M* is the list of mismatch positions. The *global* score for the candidate guide is then obtained by combining all the local scores:

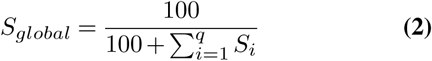

where *S*_*i*_ represents the local score between the candidate guide and off-target site *i* that partially matches the target (calculated using Eq. 1), and *q* is the number of such sites that partially matched.

Published in 2014 like Cas-OFFinder, the *mm10db* method uses a custom implementation of that score [6], in a multithreaded function called *findMismatches*. The objective of that function is to reject all guides for which the global score is below a fixed threshold *τ* = 0.75. Based on Eq. 2, *S*_*global*_ *≤ τ* is equivalent to Eq. 3.

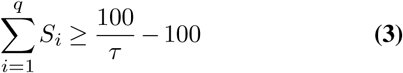

Instead of calculating an exact global score, it is therefore possible to keep track of the running sum of local scores, and terminate early as soon as a guide is guaranteed to end up being rejected.

The initial implementation of findMismatches relies on string comparisons. For this paper, we have reimplemented it using a binary encoding of the sequences, so that mismatches can be identified more efficiently.

The recently published *Crisflash* also relies on the Zhang score to evaluate guides, but without the early termination. It is reported that Crisflash can out-perform Cas-OFFinder by an order of magnitude on the human genome when more than a couple of hundred guides are supplied [7]. Crisflash also provides support for variants, which are not used here. It is also noted that Crisflash relies on a volume of memory which would not be available on a modern workstation (90 Gb), but rather a server cluster.

*FlashFry* presents a discovery approach that utilises a table of bit-encoded off-target prefixes [8]. Similar to other methods, a candidate guide is only scored on a given off-target if the number of mismatches is fewer than the specified threshold. FlashFry does not report the Zhang score, but rather the cutting-frequency determination (CFD) score [9]. The authors claim that FlashFry is able to run two to three orders of magnitude faster than Cas-OFFinder.

Other tools such as CRISPRseek [10], CasOT [11] and CRISPRitz [12] exist. However, we do not consider them here as already published results show that they perform poorly compared to others which we have included [7, 12].

## Materials and Methods

In this article, we present Crackling, a method aimed at addressing the two challenges of efficiency and specificity when designing CRISPR-Cas9 guide RNAs. To increase the efficiency of the set of guides we produce, we combined three scoring strategies. To speed up the off-target scoring, we introduce a solution for constant-time lookup of sequence neighbourhoods.

The tool is implemented in Python (version 3) for most steps, with the high-performance off-target scoring in C++. It can run on any platform. Minimal dependencies are required: Bowtie2 [13] for guide realignment onto the input genome and RNAfold [14] for secondary structure prediction. Where possible, components of the tool are parallelised for improved performance. Pre-processing is minimal, only requiring the input genome to be indexed by Bowtie2 and off-target sites to be indexed for ISSL. The ISSL index is simple to build and can be reused at any time for the genome it is constructed for.

A well-documented, Python-based configuration method is implemented. Once configured, the pipeline is called via any command line terminal using the Python v3 interpreter. Post-processing, which provides the ability to annotate guides with any genomic features which they target, is an optional final step, for which we also provide code.

### Multi-approach efficiency evaluation

We previously showed that, when tools actively filter or score guides based on their predicted efficiency, they only rarely agree with each other [1]. We also showed that this can be leveraged for consensus-based approaches that combine the output of multiple tools [2]. However, having to install and run multiple tools limits how practical such approaches can be.

Crackling directly incorporates three scoring approaches and only recommends candidate guides that have been accepted by at least two of the approaches:

1. As in CHOPCHOP [15], we are looking for the presence of a guanine at position 20 (also known as the *G20* rule).
2. We score guides based on the model from sgR-NAScorer 2.0 [16], and consider candidate guides to be accepted if their score is positive.
3. We used the filtering steps from mm10db [6] (GC content, secondary structure, etc.), and only accept candidate guides that passed all steps.

The sgRNAScorer 2.0 model is included in the Crackling code repository, in addition to the raw data and a script to retrain the model for version compatibility reasons.

### Off-target scoring using Inverted Signature Slice Lists

*Inverted Signature Slice Lists* (ISSL) can be used to rapidly perform approximate nearest neighbourhood searches in collections of locality-sensitive signatures [17]. By using fixed-length signatures as search-keys, items in a neighbourhood can be found in constant time. This approach was initially proposed as a high-performance method for searching web-scale collections of data. Given that genomic data is at a comparable scale, it provided a feasible method for evaluating CRISPR guide specificity.

The approach utilises a bit-encoded index to reduce memory and storage requirements, and to use CPU instructions most effectively. The four letters of the genomic alphabet (*A, T, C* and *G*) are two-bit encoded, hence a 20-bp genomic sequence requires 40-bits but is stored in a 64-bit word. In the context of Crackling, this 20-bp sequence represents an off-target site. For each bit-encoded signature, the critical 40-bit segment is portioned into *n* + 1 slices, where *n* is the maximum number of mismatches (here, *n* = 4 to match other methods). This slice, in combination with its position *n* is used as key within the key-value structured index. For each key, a list of off-target identifiers exists. Each identifier is also prefixed with a count of how many times the off-target site is seen across the genome. This allows the scoring method to preemptively consider all duplicates, rather than each individually, as when this would be needed with a sequential-styled approach.

Separately, when considering guide specificity, the bit-encoding process is repeated for each candidate guide. Given that *n* mismatches are allowed and there exist *n* + 1 slices, there will always be one slice shared between a candidate guide and a potential off-target site. The time complexity of this discovery step is constant given the structure of the index, and the number of potential off-target sites is less than that in other approaches (such as using a sequential search where all off-target sites are considered). Similar to the find-Mismatches approach, the candidate guide is scored against off-target sites using the Zhang score until a specified threshold is reached (Eq. 3). This approach is visualised in Figure 1.

**Fig. 1.**
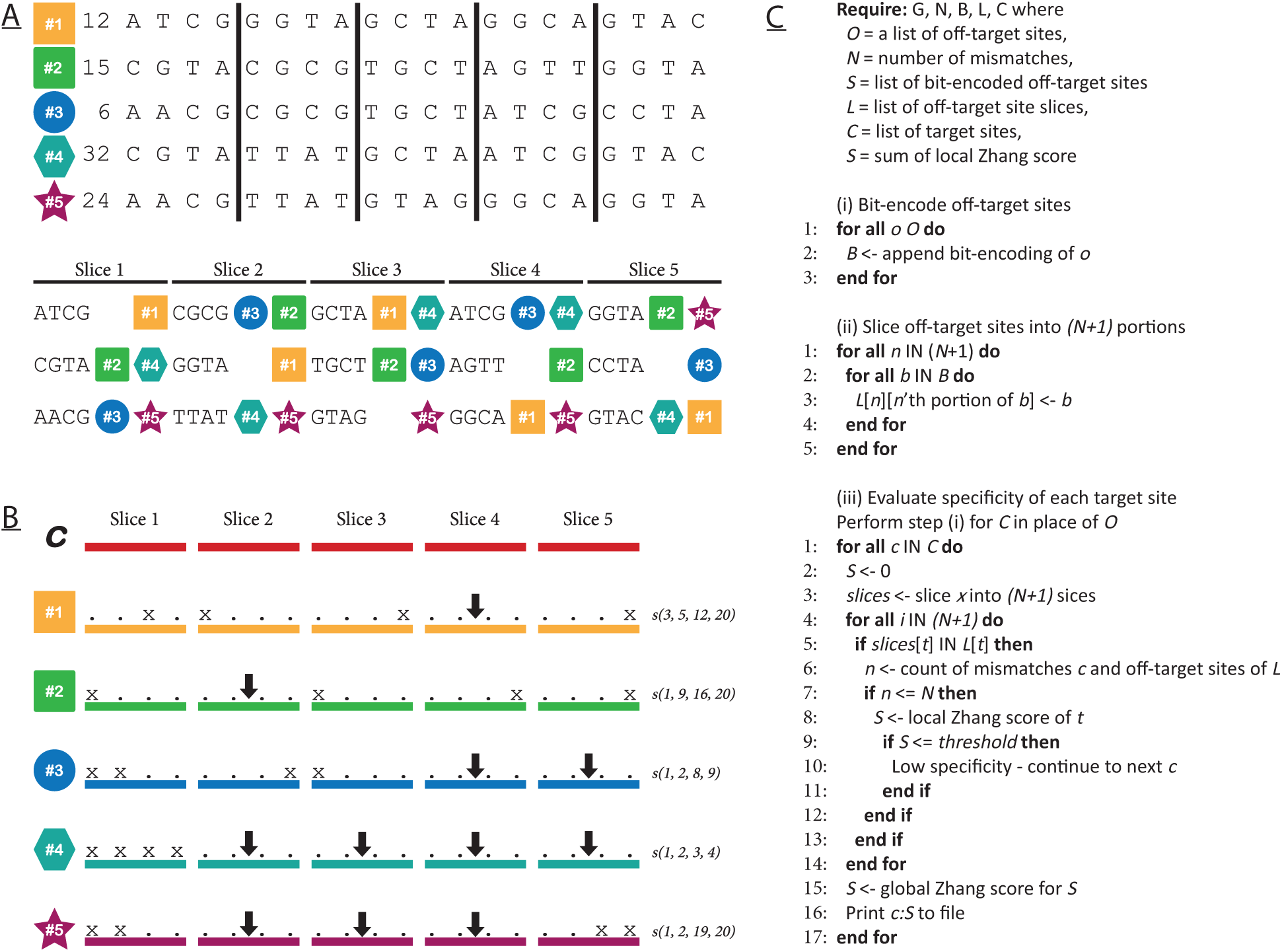
**A:** The ISSL off-targets index is a dual-key tabular data structure, constructed using locality-sensitive hashes. The key of the first dimension is the slice number. The key of the second dimension is the bit-encoded genomic sequence of the slice. The contents is the bit-encoded genomic sequence of the off-target site (depicted by the coloured shapes). **B:** Scoring a candidate guide against off-target sites. An *x* denotes a mismatch between the candidate guide and the off-target site. An arrow indicates that the given slice is common for both sequences. Each candidate guide is portioned into *n* + 1 slices (*n* = 4 in this figure). Each matching slice is used a locality-sensitive look-up key into the index. For each off-target site, the positions of mismatches are used in calculating the specificity score. If more than four mismatches exist, then the sequence is not deemed as an off-target site. An off-target may be identified for multiple slices, however ISSL strictly considers all distinct off-targets per candidate guide (e.g. off-target #3 has been identified twice but only considered for scoring once). **C:** Algorithm for ISSL

### Off-target scoring performance

We timed each of the off-target tools on genomes of increasing size, which were obtained from the National Centre for Biotechnology Information (NCBI), see Table 1. For each genome, candidate guides were extracted from the forward and reverse strands using a regular expression: [*ACG*][*AT CG*]^20^*GG* (and the complement for the reverse strand). Two editions of these target sites were created: one including the PAM sequence and one without, as this was required by the tools. This process was repeated for off-target sites, however using the following expression: [*ACG*][*AT CG*]^20^[*AG*]*G* (again, with a complementing expression for the reverse strand).

**Table 1.** The genomes used in this study. *Mb* is megabases. An asterisk indicates these genomes were only used on the high-performance machine due to memory limitations.

| Genome | Size | Off-target sites count |
| --- | --- | --- |
| *O.sativa* | 374.4 Mb | $61.7 \times 10^6$ |
| *P.hallii* | 507.4 Mb | $91.9 \times 10^6$ |
| *X.couchianus* | 685.5 Mb | $100.6 \times 10^6$ |
| *P.major* | 998.3 Mb | $173.0 \times 10^6$ |
| *C.viridis* | 1,297.0 Mb | $193.8 \times 10^6$ |
| *D.rerio* | 1,679.4 Mb | $214.8 \times 10^6$ |
| *O.mykiss* | 1,950.0 Mb | $295.9 \times 10^6$ |
| *M.musculus* | 2,730.7 Mb | $499.7 \times 10^6$ |
| *M.domestica* | 3,502.4 Mb | $570.9 \times 10^6$ |
| *T.urartu* | 4,661.6 Mb | $829.4 \times 10^6$ |
| *R.bivittatum\** | 5,178.6 Mb | $984.7 \times 10^6$ |
| *T.turgidum\** | 9,964.3 Mb | $1,763.2 \times 10^6$ |

**Table 2.** Run time for when increasing the number of candidate guides to evaluate (mm:ss). Tested on *O. sativa*.

| Dataset | ISSL | fMb | fM | Cas-OFFinder | Crisflash |
| --- | --- | --- | --- | --- | --- |
| 5k | 0:04 | 1:12 | 5:22 | 23:11 | 6:33 |
| 10k | 0:09 | 2:24 | 10:46 | 46:17 | 13:04 |
| 15k | 0:11 | 3:34 | 16:02 | 68:51 | 19:35 |
| 20k | 0:14 | 4:47 | 21:23 | 92:10 | 26:10 |
| 25k | 0:18 | 5:58 | 26:44 | 115:11 | 32:48 |

Using the candidate guides extracted for each genome, we further extract five distinct sets of 10,000 candidate guides. For *O. sativa*, we repeated this but for sets of size 5,000 to 25,000, incrementing by 5,000. For each of these, we tailored copies for Crisflash and Cas-OFFinder which required custom formats. ISSL required an index of the off-target sites. Crisflash required a single genome file, as opposed to a file for each chromosome. For this, we interspaced each chromosome with sufficient *N*’s to prevent the tool from detecting target sites which may overlap chromosomes. FlashFry required a custom index for each genome but could not produce one larger than that for *O. sativa*.

For findMismatches, findMismatchesBit, ISSL and Crisflash, we modified the source code to time the guide specificity evaluation method in each tool. Additionally, for Crisflash, we made modifications to time the construction of its tree data structure. For Cas-OFFinder, we timed it with the *time* application with microsecond precision. All preliminary tests were performed on a high-performance Linux workstation, with two 18-core Intel Xeon E5-2699 v3 (2.3 GHz) CPU’s, 512 GB RAM, 4.2 GB allocated swap space and Hewlett-Packard Enterprise MB4000GEFNA 4TB HDD. Further testing was performed on a different Linux workstation with one 8-core Intel Core i7-5960X (3.0 GHz) CPU, 32 GB RAM, 32 GB allocated swap space, and Samsung PM87 SSD.

For all tests, we used a strict 30-hour wall time, as tools slower than this would not scale to the analysis of entire genomes.

### Benchmarking guide design pipelines

We previously benchmarked leading CRISPR guide design methods using custom datasets derived from the mouse genome [1]. These datasets were of size 500,000 bp (*500k*), 1,000,000 bp (*1m*), 5,000,000 bp (*5m*) and the full 61m bp chromosome 19 (*full*). Some tools require an annotation file, and only consider guides in coding regions. To account for this we calculated the *effective base-pairs per second* (EFPS), which uses the length of the regions used for candidate extraction. Crackling considered the entire input genome, thus the genome length was used when calculating EFPS. We constructed the ISSL index for these datasets and ran identical experiments on Crackling from the original paper.

## Results

All preliminary tests were executed on a high-performance machine and followed up on a machine where the available memory is more limited. Cas-OFFinder could not be tested on the high-performance machine due to compatibility issues. For FlashFry, it was not possible to generate the required indexes for genomes larger than *O. mykiss*, and the tool was therefore not tested past this point. Generating the index for the larger genomes was attempted on both machines, with no success.

### The ISSL approach powering Crackling is the fastest off-target scoring method

Three tools (findMismatches, findMismatchesBit and ISSL) were capable of completing tests on all twelve genomes on the high-performance machine. They also completed tests on the ten genomes considered on the workstation (with the two largest being excluded due to the memory limitation of this machine). On the high-performance machine, Crisflash failed on genomes larger than *O. mykiss* due to segmentation faults. On the workstation, it saturated physical memory, causing swapping to occur.

For both machines, the run time for findMismatches and findMismatchesBit were proportional, confirming that bit-encoding alone provides a performance benefit. For example, on the high-performance machine with *C. viridis*, using bit-encoding provides a 6.6x speed-up. The results for the *T. urartu* dataset on the workstation are an outlier due to memory being swapped to disk.

ISSL was able to evaluate 10,000 candidate guides for *O. sativa* in three seconds on average; whereas, findMismatches completed analysis of the same datasets in an average of 7.5 minutes. This is a 153x speed-up. ISSL also significantly outperforms Cas-OFFinder. For instance, it completed the *T. urartu* test in 5.5 minutes on the workstation, compared to 10 hours for Cas-OFFinder. This is a 112x speed-up.

The mean run times for each test are given in Table 3, and visualised in Figure 2. ISSL is the fastest method for off-target scoring. For large genomes on low-memory machines, findMismatchesBit can also be a useful alternative, due to its lower memory footprint.

**Table 3.**
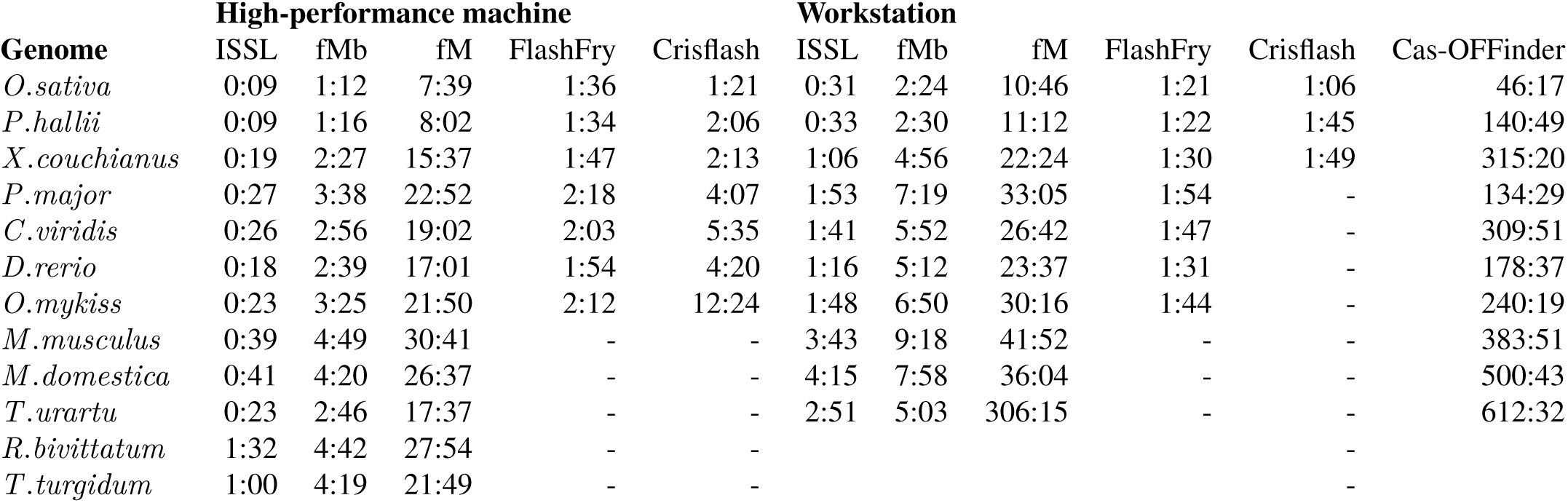
Mean run time, of 5 tests, when genome size is increased (hh:mm:ss). *-* indicates the test failed due to memory limitations. *fMb* is findMismatchesBit. *fM* is findMismatches.

**Fig. 2.**
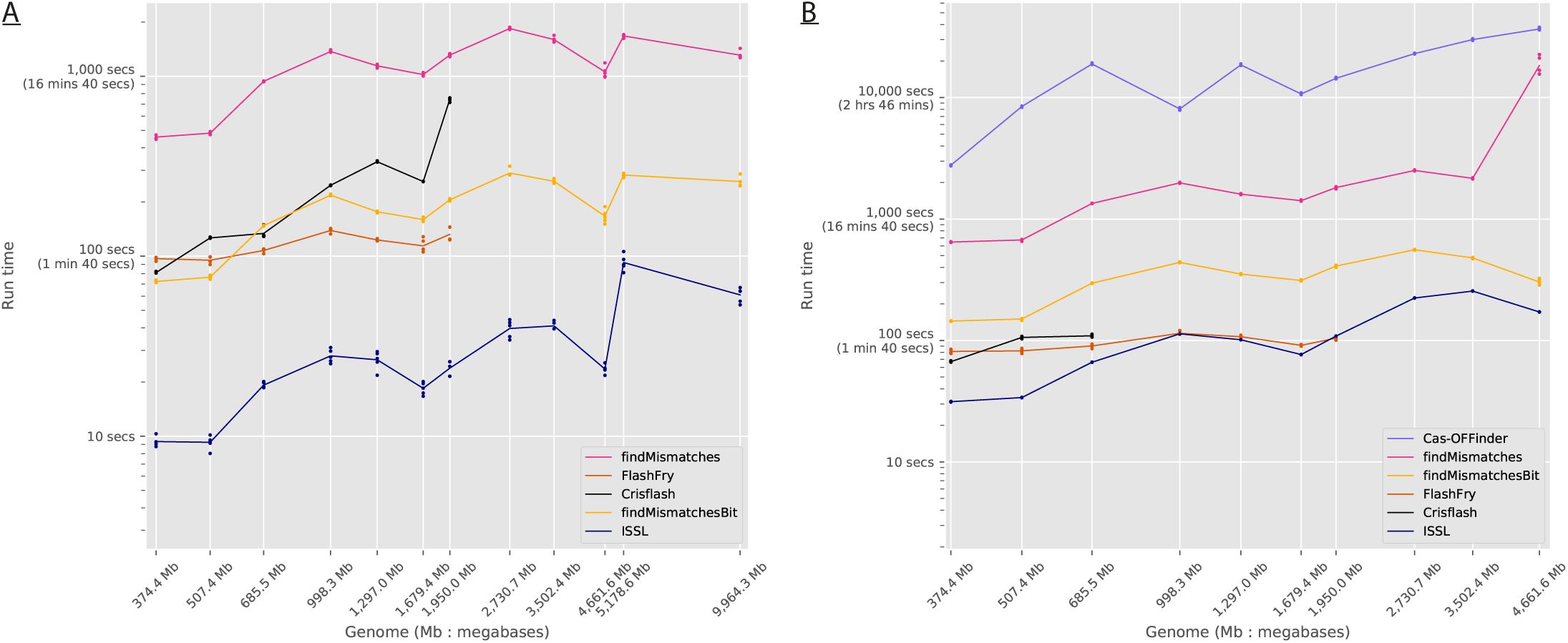
Run time on the high-performance machine (Panel A) and workstation (Panel B) for an increase in genome size. The memory requirements of Crisflash on the workstation exceeded that available and caused swapping to occur; these tests were stopped.

For ISSL and Flashfry, the supporting indexes containing off-target sites are generated in a pre-processing step that is required only once. For Crisflash, the tree data structure is constructed at run-time so we modified the source code to include a timer for this. We timed the construction of the indexes for each of these tools and report the results in Table 4, and Figures 3.

**Table 4.**
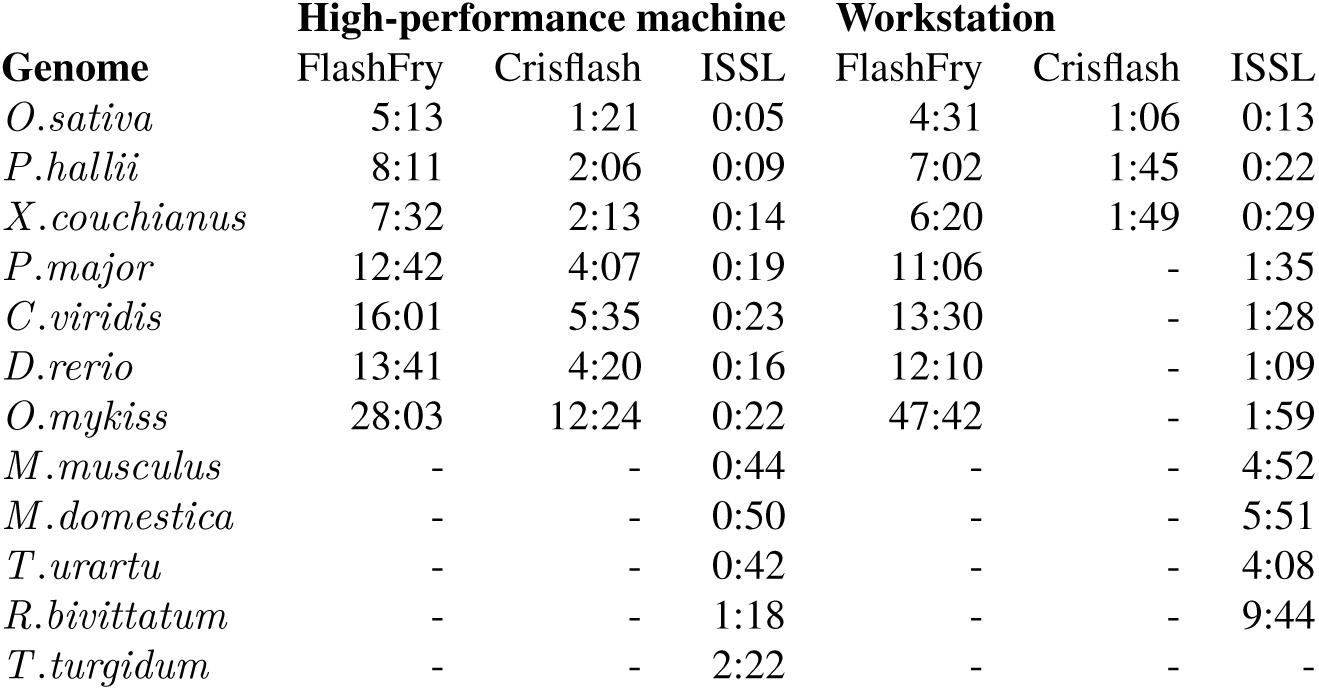
The mean run time for generating the off-targets index, of 5 tests, when genome size is increased (mm:ss). *-* indicates the test failed due to memory limitations.

| Genome | High-performance machine |  |  | Workstation |  |  |
| --- | --- | --- | --- | --- | --- | --- |
|  | FlashFry | Crisflash | ISSL | FlashFry | Crisflash | ISSL |
| *O.sativa* | 5:13 | 1:21 | 0:05 | 4:31 | 1:06 | 0:13 |
| *P.hallii* | 8:11 | 2:06 | 0:09 | 7:02 | 1:45 | 0:22 |
| *X.couchianus* | 7:32 | 2:13 | 0:14 | 6:20 | 1:49 | 0:29 |
| *P.major* | 12:42 | 4:07 | 0:19 | 11:06 | - | 1:35 |
| *C.viridis* | 16:01 | 5:35 | 0:23 | 13:30 | - | 1:28 |
| *D.rerio* | 13:41 | 4:20 | 0:16 | 12:10 | - | 1:09 |
| *O.mykiss* | 28:03 | 12:24 | 0:22 | 47:42 | - | 1:59 |
| *M.musculus* | - | - | 0:44 | - | - | 4:52 |
| *M.domestica* | - | - | 0:50 | - | - | 5:51 |
| *T.urartu* | - | - | 0:42 | - | - | 4:08 |
| *R.bivittatum* | - | - | 1:18 | - | - | 9:44 |
| *T.turgidum* | - | - | 2:22 | - | - | - |

**Fig. 3.**
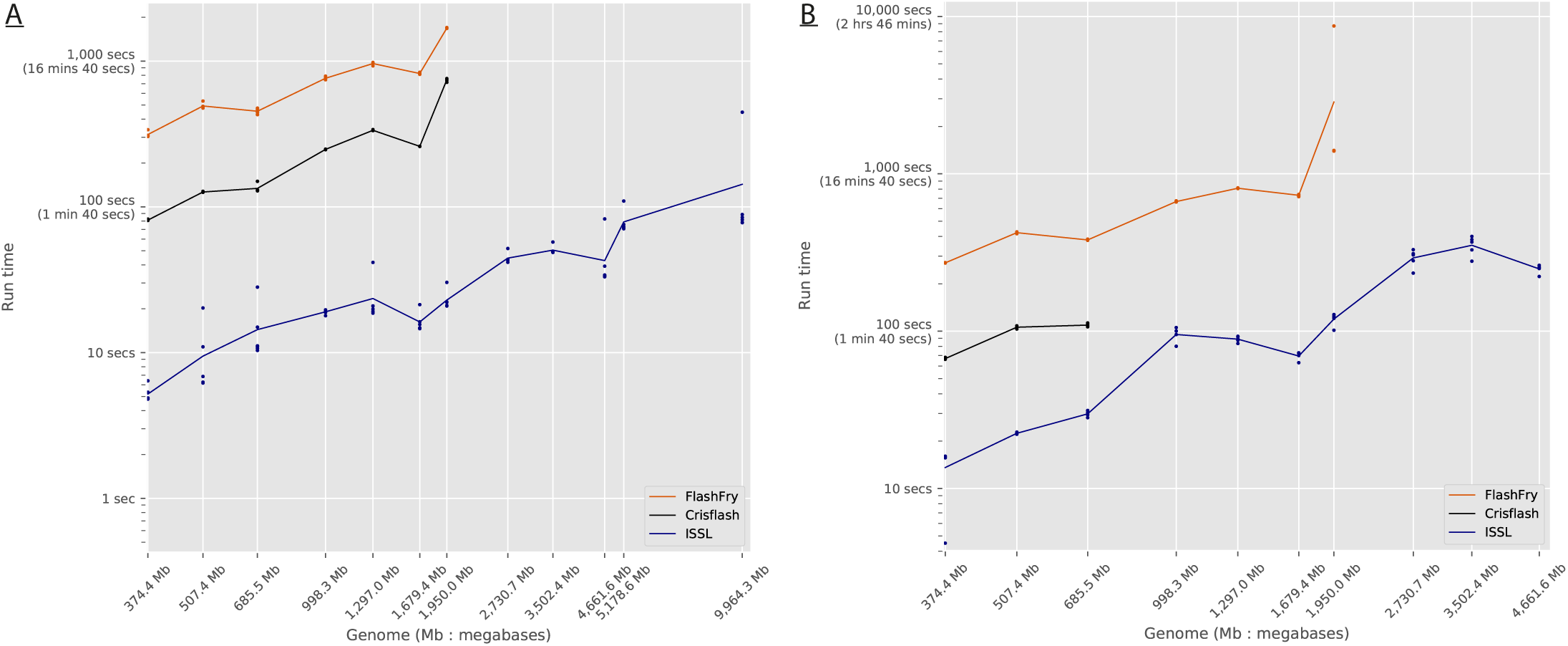
Run time for generating off-targets indexes on the high-performance machine (Panel A) and workstation (Panel B).

### Impact of the number of targets

We ran each tool on the *O. sativa* genome and varied the number of candidate guides to be evaluated. The results are shown in Table **??** for mean run times over five tests.

ISSL completed all tests in under 25 seconds, whereas Cas-OFFinder completed the test on the smallest sized dataset in 23 minutes and largest sized dataset in 2 hours. Here, ISSL performed up to 300x faster than Cas-OFFinder, and at least an order of magnitude faster than the next tool.

### Crackling is a fast method to produce efficient guides

We constructed the ISSL index for these datasets and ran identical experiments on Crackling from the original paper. Crackling was able to complete five repeat tests on each genome without failure. The average EFPS for the datasets were:

- *500k*: 11,027 EFPS
- *1m*: 12,459 EFPS
- *5m*: 21,643 EFPS
- *full*: 32,625 EFPS

This is also visualised in Figure 4. In the benchmark, we used this Figure to define low-, medium- and high-performance groups. Crackling enters the high-performance group, and is the only method in that group to correctly filter guides (Cas-Finder and CRISPR-ERA provide a score, but it did not prove to be informative). Compared to other tools that filter guides, Crackling is at least an order of magnitude faster.

**Fig. 4.**
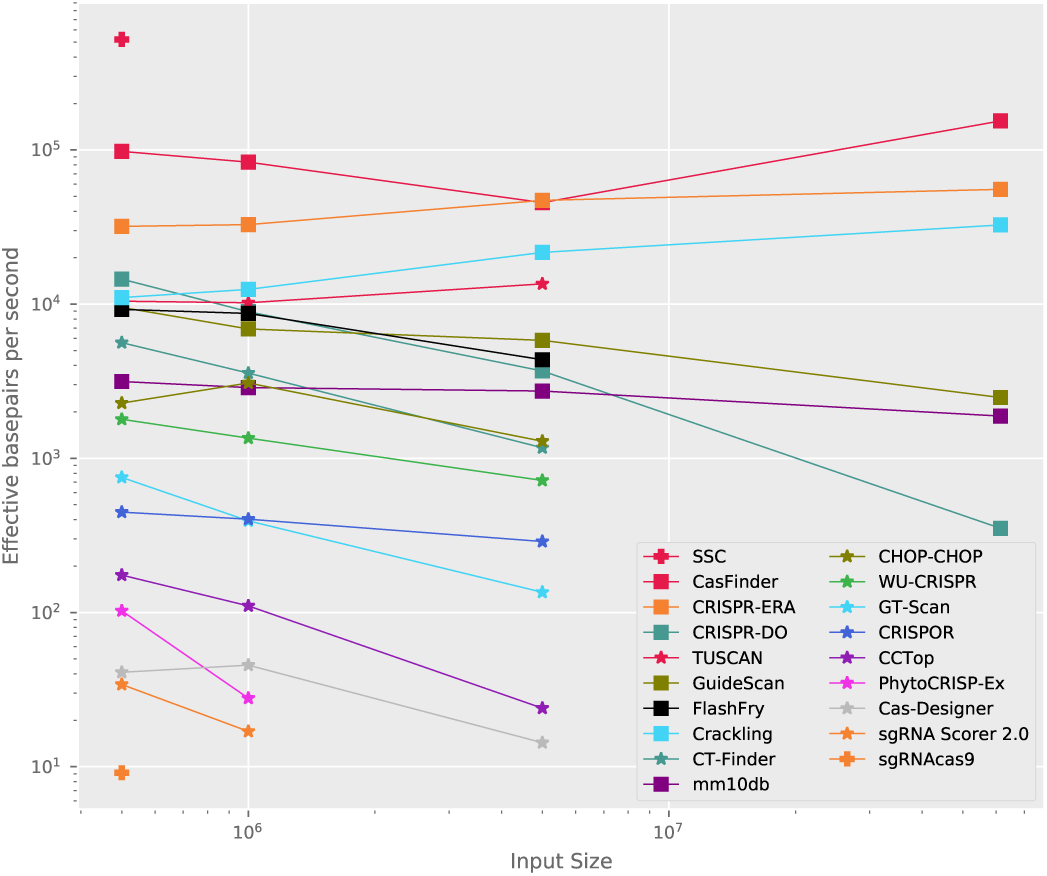
Run time on mouse chromosome 19 datasets. This plot is extended from [1] to include Crackling.

The quality of the guides produced by Crackling is also the highest amongst the tools reviewed. Table 5 describes the precision of each tool on two experimentally validated datasets, *Wang* [18] and *Doench* [19]. As seen in Figure 5A for the *Doench* dataset, Crackling correctly selected many guides that are *efficient* and very few that are *inefficient*, and again for the *Doench* dataset in Figure 5B. Notably, Crackling, by default, requires a candidate guide to be accepted by two of the three scoring approaches in order to be selected. When increasing this to require all three approaches to agree, the precision of Crackling will increase (to 91.2% and 48.5% on the *Wang* and *Doench* datasets, respectively), but the recall drops to values that are only acceptable when a low number of candidates is needed (7.1% and 8.9%, respectively).

**Table 5.**
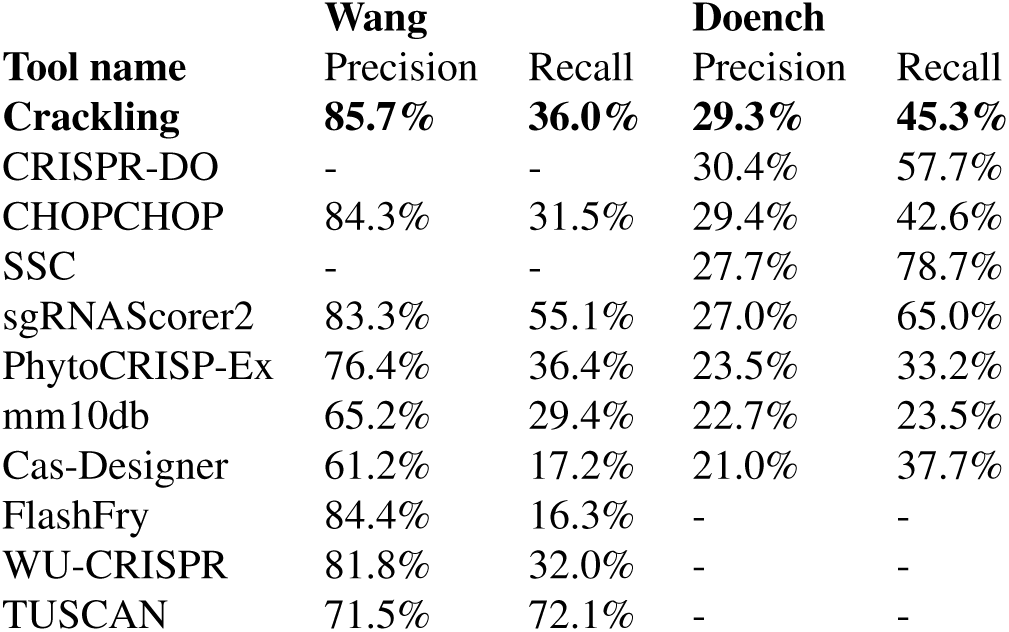
Precision of tools on experimentally validated datasets.*-* indicates the tool was not tested on this dataset as it was used for training. Adapted from [2].

| Tool name | Wang |  | Doench |  |
| --- | --- | --- | --- | --- |
|  | Precision | Recall | Precision | Recall |
| **Crackling** | **85.7%** | **36.0%** | **29.3%** | **45.3%** |
| CRISPR-DO | - | - | 30.4% | 57.7% |
| CHOPCHOP | 84.3% | 31.5% | 29.4% | 42.6% |
| SSC | - | - | 27.7% | 78.7% |
| sgRNAScorer2 | 83.3% | 55.1% | 27.0% | 65.0% |
| PhytoCRISP-Ex | 76.4% | 36.4% | 23.5% | 33.2% |
| mm10db | 65.2% | 29.4% | 22.7% | 23.5% |
| Cas-Designer | 61.2% | 17.2% | 21.0% | 37.7% |
| FlashFry | 84.4% | 16.3% | - | - |
| WU-CRISPR | 81.8% | 32.0% | - | - |
| TUSCAN | 71.5% | 72.1% | - | - |

**Fig. 5.**
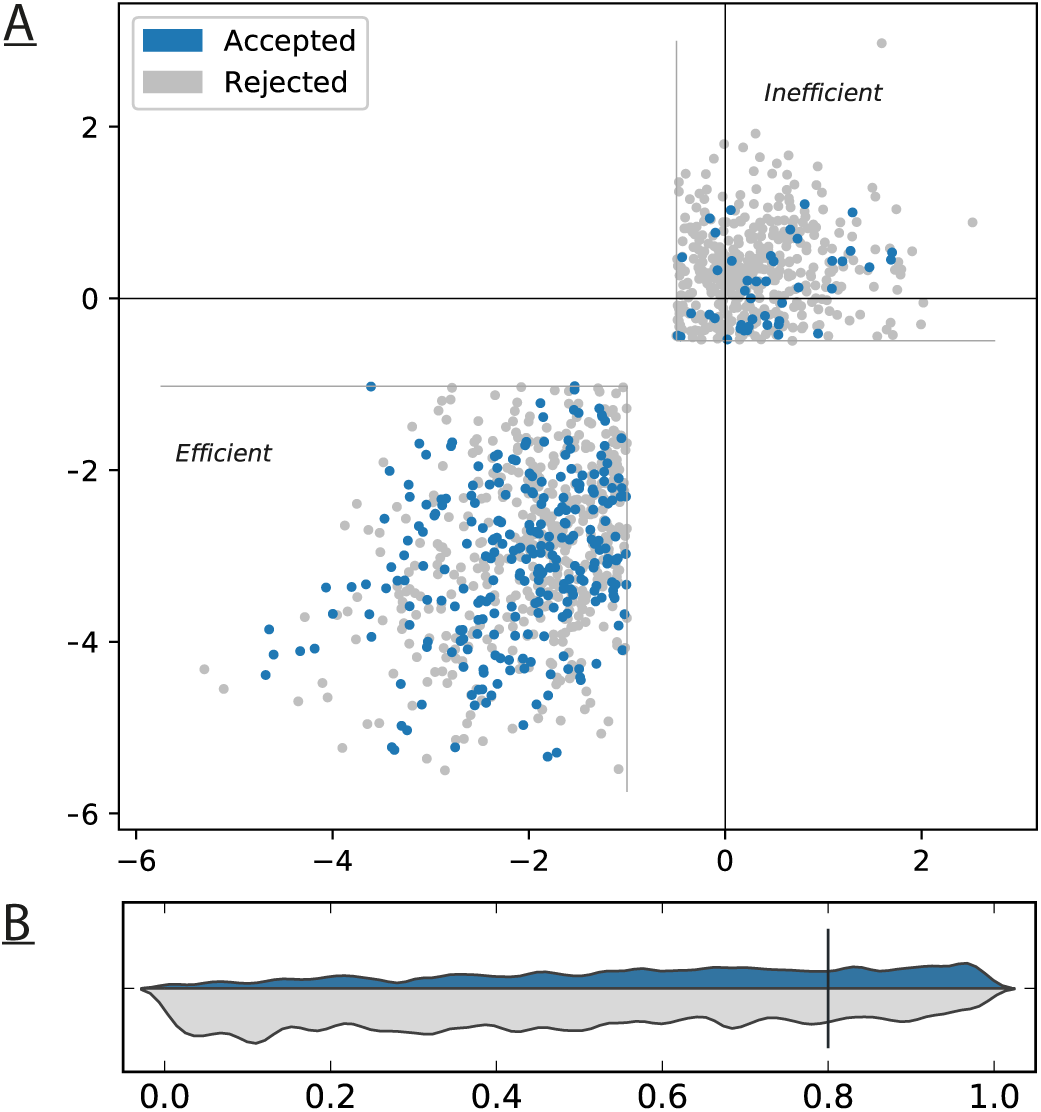
Results on the Doench dataset. As per [2], “The blue distribution shows the number of guides accepted, and the grey distribution shows the number of guides rejected. The vertical marker at 0.8 shows the threshold used to determine efficiency; guides with a gene rank score greater than this were deemed experimentally efficient”.

## Discussion

One of the main challenges of experiments using CRISPR-Cas9 systems is the design of suitable guide RNAs. It is crucial, but not trivial, to ensure that these guides will be efficient, but also specific enough not to lead to off-target modifications. There was no tool that could meet both objectives while remaining practical to use on large genomes. In this paper, we presented Crackling, a new tool for whole-genome identification of suitable CRISPR targets. Crackling is powered by a new high-performance off-target scoring approach based on Inverted Signature Slice Lists (ISSL) and utilises our previously published consensus approach for improving the precision when predicting guide efficacy.

When executing ISSL on a set of genomes using a high-performance machine, ISSL performs approximately an order of magnitude faster than the next best performing tool (findMismatchesBit), and up to two orders of magnitude faster than the worst performing tool (findMismatches).

We ran a number of off-target scoring tools on the *O. sativa* genome and varied the number of candidate guides to be evaluated. It would be expected that runtime is a linear function of input size, seeing that each candidate guide is evaluated individually. For all tools, we found that this is true.

Overall, our findings are that Cas-OFFinder is the poorest performing tool; findMismatches and findMismatchesBit are proportional; ISSL is the best performing, followed by Crisflash and finally Flashfry.

The precision of Crackling is greater than any tool taken individually. This is achieved via a consensus approach which only accepts guides that have been predicted as efficient by at least two of three scoring methods. Further gains in precision can be achieved by changing this to be a consensus of all three scoring methods.

Taken together, all these results show that Crackling is able to produce high quality results in a relatively short period of time. It is able to perform better than any tool on both critical properties (specificity and efficiency) that are considered when evaluating CRISPR guides.

We showed that it provides the fastest way to calculate the off-target risk, by using binary encoding and the ISSL method to identify closely related throughout the genome. This was confirmed on twelve genomes of increasing length. We also showed that by combining the results of multiple scoring approaches, we can maximise the efficiency of the guides being designed. On experimental data, the set of guides it selects are better than those produced by existing tools.

Overall, this makes Crackling a faster and better method to design guide RNAs at scale.

## Acknowledgements

We would like to thank the Big Data lab at the Queensland University of Technology for providing access to their high-performance machine, and the authors of the other tools for making their source code available.

